# An Efficient Workflow for CHO Cell Genome Engineering with OpenCRISPR-1

**DOI:** 10.64898/2025.12.18.695036

**Authors:** Divya Kolakada, Ramya Ankala, Joanna Donatelli, Derek Smith

**Affiliations:** Gilead Sciences, Foster City CA

## Abstract

The effective titer and quality of biopharmaceutical products can be enhanced by genome engineering of producer cell lines; however, licensing constraints often limit nuclease utility. Here, we validate OpenCRISPR-1 in CHO cells by achieving ≥70% INDEL efficiency across multiple genes. We demonstrate biallelic *Fut8* knockout in monoclonal cell lines with 31% efficiency and quadruplex knockout of lipases at 7% efficiency using a 39-day workflow. This work highlights the potential applications for democratized nucleases in host cell engineering.

## Introduction

The past decade witnessed the discovery of diverse genome engineering tools including CRISPR-Cas^1^, adenine and cytosine base editors^2,3^, prime editors^4^, and bridge RNAs^5^. However, commercial application of these cutting-edge technologies to manufacturing cell lines has remained limited by various constraints. This has driven continued use of established but less efficient tools like zinc finger nucleases (ZFNs) and transcription activator-like effector nucleases (TALENs) to engineer cell lines for commercial manufacturing applications.^6^

Recent advances in generative protein language models have enabled the computational prediction of new-to-nature nucleases.^7^ OpenCRISPR-1 is an exemplary outcome of this modeling – both as a specific and efficient nuclease, as well as, an open-access tool for both research and commercial applications.^7^ This first-of-a-kind nuclease has shown comparable activity to SpCas9 in human cell lines; however, remains untested in Chinese Hamster Ovary (CHO) cell lines, a workhorse for commercial biologics production. Here, we demonstrate OpenCRISPR-1 activity across 10 different genomic targets in CHOZN cells. Further, we open-access an efficient workflow for generating clonal single-gene and multiplexed gene knockout in CHO cell lines.

## Evaluating Democratized Nucleases in CHO Cell Lines

One bottleneck to achieving efficient nuclease activity in mammalian cell lines is delivery and transport of nuclease machinery into the cell nucleus. To tease apart transfection efficiency from nuclease activity, we co-transfected CHOZN cells with an mRNA encoding OpenCRISPR-1 nuclease with nuclear localization sequences on both the N- and C-termini and sgRNAs targeting the genomic hotspot *LOC100768845*^8^ using either electroporation or lipofectamine (Supplemental Figure 1A). Electroporation achieved INDEL generation efficiencies exceeding 90% for multiple guide RNAs (Figure 1A). In addition to paired-end next generation sequencing, we verified INDEL generation by deconvolution of low-cost Sanger sequencing reactions using Tracking of Indels by Decomposition (TIDE) software (Supplemental Figure 1B).^9^ Interestingly, we observed higher nuclease activity and more functional sgRNAs targeting *LOC100768845* for OpenCRISPR-1 nuclease than MAD7 nuclease (Supplemental Figure 1C). Selecting the highest efficiency guide sequence for each nuclease, we next demonstrated on-target single-copy knock-in of a 2.2 kb landing pad in clonal cell lines using junction ddPCR then confirmatory targeted locus amplification sequencing (Supplemental Figure 1D). Collectively, this dataset demonstrates for the first time that a zero-cost AI-generated nuclease can perform knock-in and knockout across a range of genomic sites in a cell line relevant to biopharmaceutical manufacturing.

**Figure 1.**
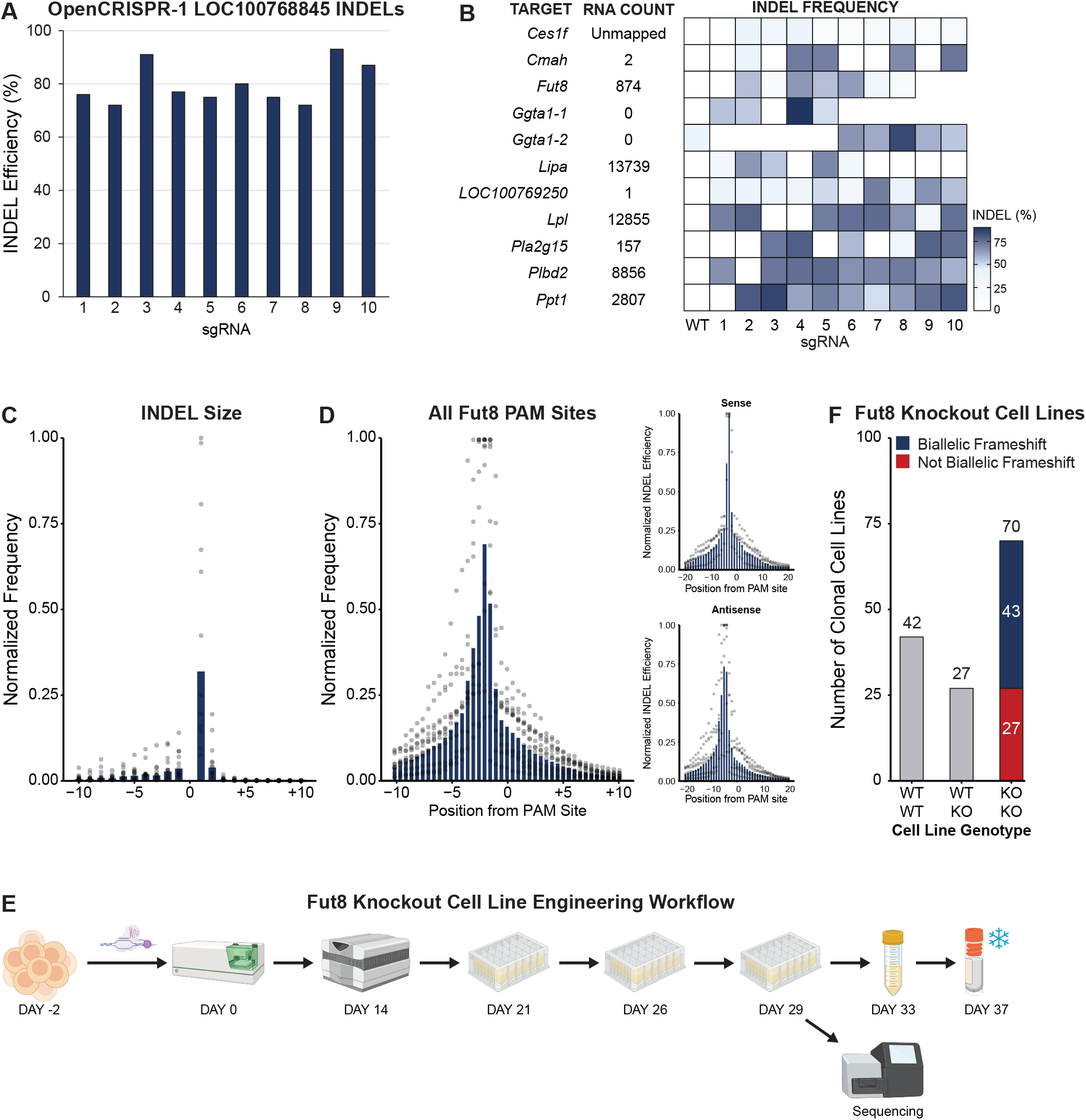
OpenCRISPR-1 Nuclease Activity and Biallelic Knockout of *Fut8* in Clonal Cell Lines. A) OpenCRISPR-1 INDEL frequency at LOC100768845. B) RNA-seq counts for ten CHO genes and the corresponding INDEL frequency for each target gene generated by OpenCRISPR-1 sgRNAs. C) The average INDEL size generated by OpenCRISPR-1 nuclease. D) INDEL position relative to the OpenCRISPR-1 PAM sequence for *Fut8* sgRNAs with subplot distributions based on location of PAM sequence on sense or antisense DNA strand. E) Clonal cell line generation workflow for *Fut8* knockout cell lines. F) Number and genotype of clonal cell lines generated by OpenCRISPR-1 knockout of *Fut8*.

## Biallelic *Fut8* Knockout using OpenCRISPR-1 Nuclease

One strategy to improve product quality and yield from producer CHO cell lines is host genome engineering. Given our robust technical successes in CHO cells and the open-access spirit of OpenCRISPR-1, we targeted a broad range of genes with the potential to improve antibody product quality including *Ces1f, Cmah, Fut8, Ggta1, Lipa, LOC100769250, Lpl, Pla2g15, Plbd2*, and *Ppt1* (Figure 1B). A screen of ten guide RNA sequences for each target gene consistently yielded 70-90% INDEL efficiency with exception of *Ces1f* at 39%. Importantly, we also evaluated published Cas9 sgRNA sequences which demonstrated high-efficiency gene editing and direct compatibility of OpenCRISPR-1 with Cas9 sgRNAs (Supplemental Figure 1E).^10-12^

To better understand the mechanism OpenCRISPR-1 uses to create double-stranded DNA breaks, we analyzed INDEL profiles for *Fut8*-targeting sgRNAs. Most often, OpenCRISPR-1 created a single nucleotide insertion four nucleotides upstream of the PAM site, indicative of a cut site consistent with Cas9 nuclease (Figure 1C, D). Interestingly, positional INDEL frequency profiles differed between the sense and antisense DNA strands with the highest INDEL frequency for the sense strand being three nucleotides upstream of the PAM and either four or five nucleotides upstream of the antisense PAM (Figure 1D). This positional effect was consistent across PAM sequences (Supplemental Figure 2).

Given the significant impact of fucosylation on antibody-dependent cellular cytotoxicity products^13-15^, we used OpenCRISPR-1 nuclease and sgRNA 6 to target exon 11 of *Fut8* to disrupt protein expression. Single cell cloning of pools was performed 48 hours post-electroporation and 139 monoclonal cell lines scaled up for next-generation sequencing and cryopreservation (Figure 1E). Targeted amplification and paired-end sequencing of *Fut8* exon 11 identified 42 unmodified (30%), 27 heterozygous cell lines (19%), and 70 cell lines with biallelic mutation (51%) (Figure 1F). Translating the disrupted *Fut8* DNA sequences into amino acid sequences, we observed 43 cell lines with biallelic frameshift. By targeting the catalytic site of this fucosyltransferase and selecting only cell lines with biallelic frameshift mutations, we achieved approximately 31% efficiency in OpenCRISPR-targeted knockout. For the first time, we demonstrate high-efficiency biallelic knockout of a gene critical to antibody product mechanism-of-action using an open-access, no cost nuclease.

## Quadruplex Knockout of Polysorbate Degrading Lipases

We developed a 36-day clonal cell line generation workflow and derived CHO cell line 1000-3310, a high-productivity subclone of the CHOZN cell line.^16^ Macroscopic visualization of the host genome by karyotyping revealed significant divergence between host cell lines, as well as sister chromosomes of each cell line (Figure 2A). Further, transcriptome profiling of 1000-3310 and the parental CHOZN host by mRNA sequencing indicates broad divergence in gene expression profiles between host cell lines (Figure 2B). Interestingly, multiple glycosylation-related host genes were not expressed in CHOZN cell lines; however, multiple lipases implicated in polysorbate degradation were robustly expressed.^17^ For this reason, we co-transfected cells with sgRNAs targeting *Lipa, Lpl, Pla2g15*, and *Plbd2*. Of the 1637 clonal cell lines printed, only 398 cell lines (24%) survived into colony formation suggesting a combination of two or more of these lipases could be important to cell survival during the stress of single cell cloning and expansion. Cell lines were scaled up, sampled for high-throughput ddPCR-based drop-off assays, and quadruplex knockout assessed in a subset of clonal cell lines using targeted amplification and sequencing of each targeted exon (Figure 2C). In an effort to rapidly screen all 398 cell lines for biallelic mutations across multiple genomic targets, we developed ddPCR-based drop off assays against OpenCRISPR-1 targeted sites in *Lipa, Lpl, Pla2g15*, and *Plbd2*.^18^ We observed high frequencies of biallelic INDEL generation at both *Pla2g15* (78.4%) and *Lipa* (68.1%), while 51.6% of all clonal cell lines had biallelic mutations in both genes. We observed lower biallelic INDEL generation frequencies in *Lpl* resulting in 50 cell lines (12.6%) with bialleic mutations in *Pla2g15, Lipa*, and *Lpl*. In total, we identified 28 clonal cell lines (7%) with biallelic mutations in all four lipase genes (Figure 2C). Of these 28 cell lines, we confirmed two encoded protein disrupting frame-shift mutations across all four genes *Lipa, Lpl, Pla2g15*, and *Plbd2*. For the first time, we demonstrate the ability to generate clonal CHO cell lines with multiplexed biallelic gene knockout using OpenCRISPR-1 nuclease.

**Figure 2.**
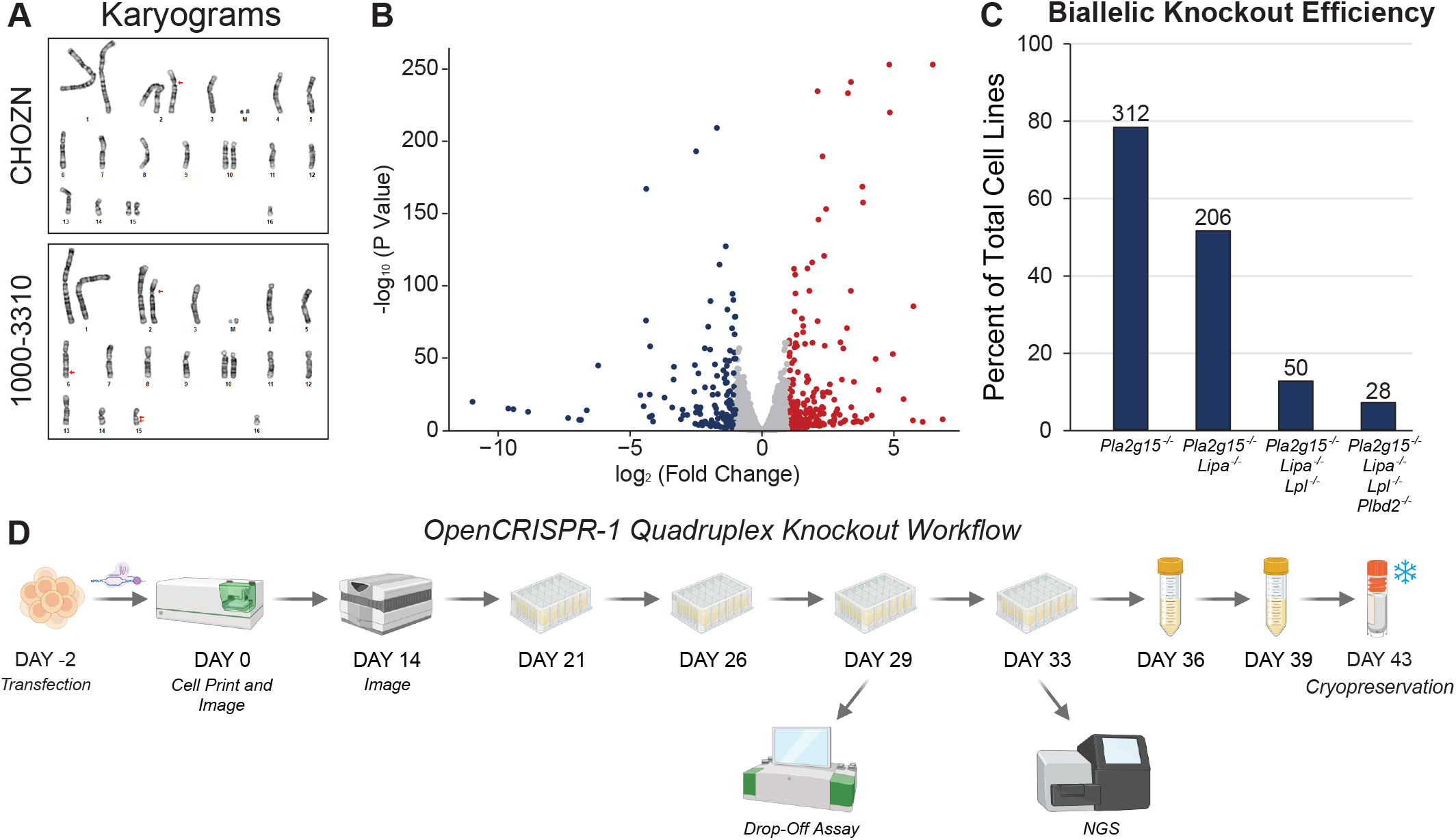
Quadruplex Knockout of Polysorbate Degrading Lipases. A) Karyogram of CHO cell lines utilized to screen sgRNAs for OpenCRISPR-1 nuclease. B) mRNA-seq profiling demonstrates statistically significant differences in gene expression between the CHOZN cell line and cell line 1000-3310, a subclone of the CHOZN parental cell line. C) The number and percent of total of each genotype of clonal cell line generated by OpenCRISPR-1 knockout of lipase genes. D) Clonal cell line generation workflow for lipase knockout cell lines.

## Methods

### Cell Culture

CHOZN GS-/-CHO cells were cultured in EX-CELL CD CHO Fusion Cell Culture Growth Medium supplemented with 3% (v/v) GlutaMAX. Suspension cell culture was performed at 37°C and 125 rpm in a humidified incubator with 5% CO_2_. Cells were passaged twice weekly, seeded first at 0.3E6 cells/mL for 3 day culture then seeded at 0.2E6 cells/mL for 4 day culture.

### OpenCRISPR-1 Genome Engineering

2E6 cells were electroporated with 2 µg OpenCRISPR-1 mRNA and 4 µg sgRNA in Genome Editing Buffer using a Neon NxT electroporation system (pulse voltage: 1650 V, pulse width: 10 ms, pulse number: 3). Landing pad knock-in experiments used 4 µg nuclease mRNA, 5 µg sgRNA, and 0.5 pmol of donor DNA.

### Single Cell Deposition and Imaging

The viable cell density of CHOZN GS CHO cells was quantified using ViCell Blu then 1E6 cells collected by centrifugation (4 minutes, 4,000 rpm, room temperature). Spent medium was aspirated and the cells were resuspended in 1 mL PBS. One vial of CellTrace Violet dye was resuspended in 20 μl DMSO then 1 μl resuspended dye added to the cell suspension. Cells were incubated for 45 minutes at 37°C and 125 rpm in a humidified incubator with 5% CO_2_. Cells were collected by centrifugation (4 minutes, 4,000 rpm, room temperature), staining solution aspirated, and cells resuspended in 1 mL PBS. Single cell cloning medium consisting of EX-CELL CHO Cloning Medium, 20% (v/v) filtered spent medium from CHOZN GS CHO cells, 2.5% (v/v) ClonaCell CHO ACF Supplement, and 3% (v/v) GlutaMAX was filtered through a 0.2 μm PES membrane. Next, 200 μl of filtered medium was added to each well of a 96-well plate and the plate incubated at 37°C until single cell deposition using an F.SIGHT 2.0. Following cell deposition, 10 μl of cell suspension was transferred to well C4 as a focus control then each 96-well plate was centrifuged (1 minute, 1,000 rpm, room temperature). A Celigo plate imager was used to acquire brightfield and fluorescence images for each well of each 96-well plate. Plates were incubated at 37°C for 9 or 14 days then, after colony outgrowth, a brightfield image of each well was captured. To each well of the 96-well plate, 100 μl EX-CELL CD CHO Fusion Cell Culture Growth Medium containing 3% (v/v) GlutaMAX was added and cells were triturated three times to disaggregate clumped cells. Images were processed for monoclonal occupancy and outgrowth to generate a hitpick list of clonal cell lines originating only from a single cell with a colony with area >5% of the well.

### Clonal Cell Line Culture and Scale Up

Clonal cell lines were hitpicked by transferring 200 μl of source cell suspension into 24-well deep-well plates containing 1 mL of warm EX-CELL CD CHO Fusion Cell Culture Growth Medium containing 3% (v/v) GlutaMAX. Cells were incubated 5 days at 37°C and 125 rpm in a humidified incubator with 5% CO_2_. The viable cell density of each clonal cell line was quantified using ViCell Blu then a volume corresponding to 0.3E6 cells transferred from the source 24-well plate into a destination 24-well deep-well plate containing (1.2 – volume of cells) mL warm EX-CELL CD CHO Fusion Cell Culture Growth Medium containing 3% (v/v) GlutaMAX. Cell passage in 24-well deep-well plates continued for three passages: first (3 days, 0.3E6 cells/mL in 1.2 mL), second (4 days, 0.3E6 cells/mL in 3 mL), and third (3 days, 0.3E6 cells/mL in 3mL). Clonal cell lines were scaled up using tubespin into 10 mL at 200 rpm. Cells were cryopreserved using a VIAFreeze Quad controlled rate freezer in EX-CELL CD CHO Fusion Cell Culture Growth Medium containing 3% (v/v) GlutaMAX and 7.5% (v/v) DMSO.

### Digital Droplet PCR Drop-Off Assay

Primer–probe mixes (20X) for *Lipa, Lpl*, and *Pla2g15* were designed using the Bio-Rad Genome Edit Detection tool. Primers, HEX drop-off probe, and FAM reference probe were designed in-house for *Pldb2* and synthesized by Integrated DNA Technologies. Genomic DNA was extracted from clonal cell lines using Lucigen Quick Extract Buffer. A ddPCR reaction mixture containing ddPCR Supermix for Probes (no dUTP), 5-125 ng diluted genomic DNA, 1 µl primer-probe 20X mix (900 nM primers and 250 nM probes), and 5 units HindIII restriction enzyme was added to nuclease-free water to 20 µL total volume. The reaction mix was loaded into a GC36 cartridge, heat-sealed, centrifuged, and processed on a QX ONE system for droplet generation followed by thermal cycling (55°C melting temperature and 5 minute extension) and droplet reading. Droplets were analyzed based on fluorescence amplitude and subsequent fractional abundance calculations were performed using QX ONE Software v1.4 Standard Edition.

### Sequencing Methods

Genomic DNA extraction, library construction, DNA sequencing, and bioinformatic analysis of INDEL frequencies for *Ces1f, Fut8, Lipa*, and *Lpl* genes was performed by Azenta Life Sciences while INDEL frequencies for the genes *Ggta1, Cmah, Plbd2, Pla2g15, Ppt1, LOC100769250*, as well as, *Fut8* clone variant identification and quadruplex lipase knockout variants were performed by Gilead Sciences. Genomic DNA was extracted using either QuickExtract Solution at 65°C for 6 minutes followed by 98°C for 2 minutes or using a Qiagen DNeasy Blood & Tissue Kit. Sequencing libraries were prepared using a two-step PCR workflow. The first PCR amplified the target region from genomic DNA using locus-specific primers with Illumina overhang adapters (Supplemental Table 1) followed by purification with magnetic beads. Indices and Illumina sequencing adapters were added using limited-cycle PCR. *Ggta1* Region 1 was the only exception to this workflow requiring nested PCR to overcome hard to amplify GC rich regions. Libraries size was analyzed using TapeStation and quantified using Qubit 4.0 Fluorometer and quantitative PCR. Equimolar libraries were pooled and sequenced on an Illumina miSeq using 2□×□300 bp paired-end configuration. Image analysis and base calling were performed using miSeq Control Software. For TIDE analysis, genomic DNA was extracted and target sequences amplified by PCR. PCR products were visualized by using TapeStation and quantified using Qubit 4.0 Fluorometer then Sanger sequenced. Sequence traces were analyzed using TIDE version 5.0.5.

## Data Analysis

Sequence analysis for *Ces1f, Fut8, Lipa*, and *Lpl* genes was performed by Azenta Life Sciences wherein raw FASTQ files were trimmed to remove low-quality reads using Sickle. Paired-end reads were merged using PANDAseq. The merged reads were aligned to the reference sequence using BWA followed by a proprietary variant detection analysis. Sequence analysis for *Ggta1, Cmah, Plbd2, Pla2g15, Ppt1, LOC100769250, and Fut8* clonal knockout cell lines cell lines was performed by merging paired-end reads and removal of low-quality reads by filtering using SeqPrep. CRISPResso2 was used to perform variant detection analysis for clone characterization and assess sgRNA cutting efficiencies for each gene. Workflow images were generated using BioRender.

