## Supplemental Figure 1 for "An Efficient Workflow for CHO Cell Genome Engineering with OpenCRISPR-1"

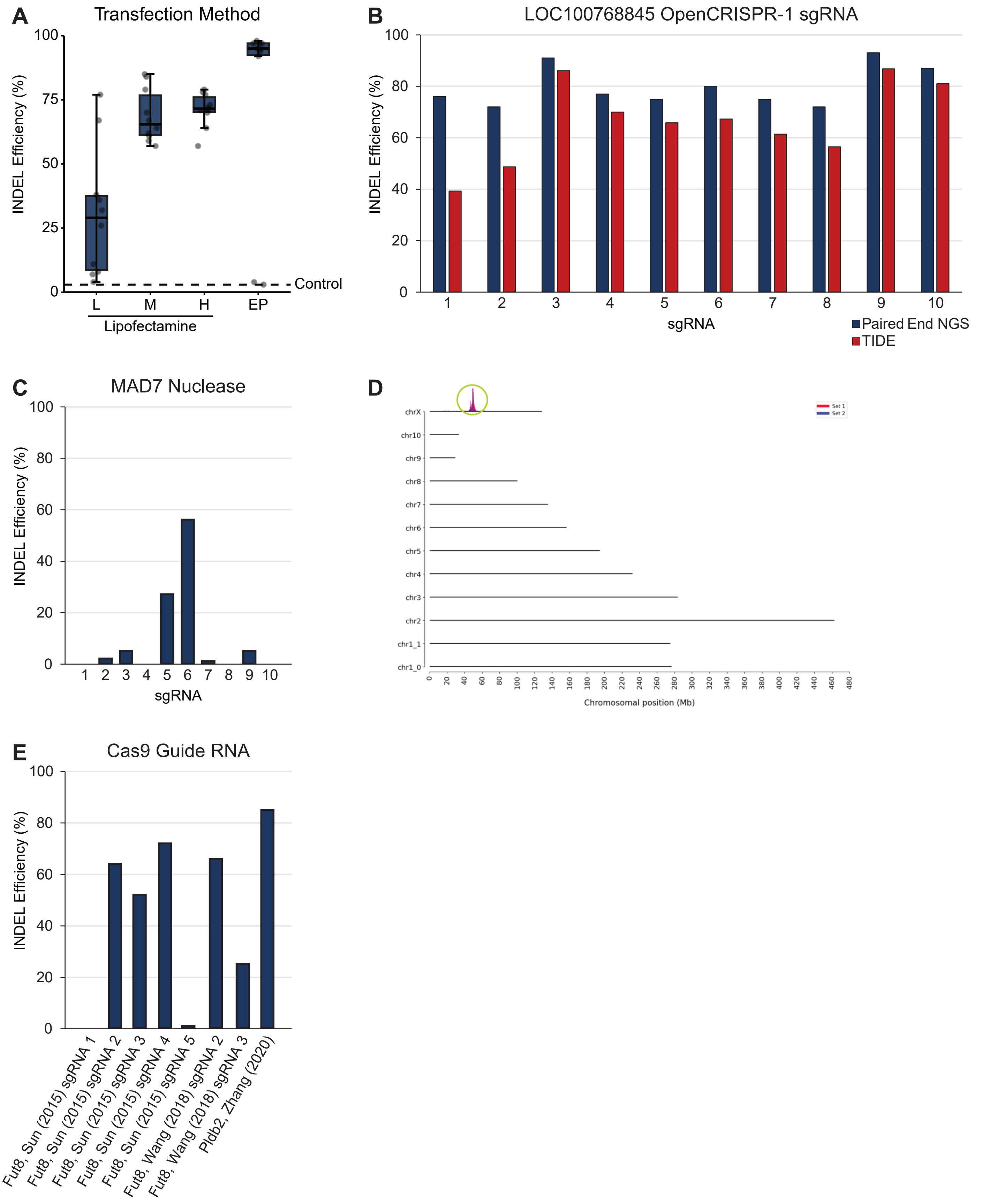


**Supplemental Figure 1. Functional Activity of Democratized Nucleases**

A) The rate of INDEL generation for cells transfected with lipofectamine using L (1 µg mRNA and 2 µg sgRNA), M (2 µg mRNA and 4 µg sgRNA), and H (4 µg mRNA and 8 µg sgRNA) relative to electroporation with a Neon NxT using M (2 µg mRNA and 4 µg sgRNA). B) Comparison of identified INDEL frequency using miSeq libraries versus Sanger sequencing and TIDE. C) MAD7 nuclease INDEL frequency at LOC100768845. D) Targeted locus amplification sequencing confirming on-target, single-copy integration of a 2.2kb landing pad at LOC100768845. E) Confirmation of OpenCRISPR-1 activity with Cas9-compatible guide RNA sequences.
