## Supplemental Figure 2 for "An Efficient Workflow for CHO Cell Genome Engineering with OpenCRISPR-1"

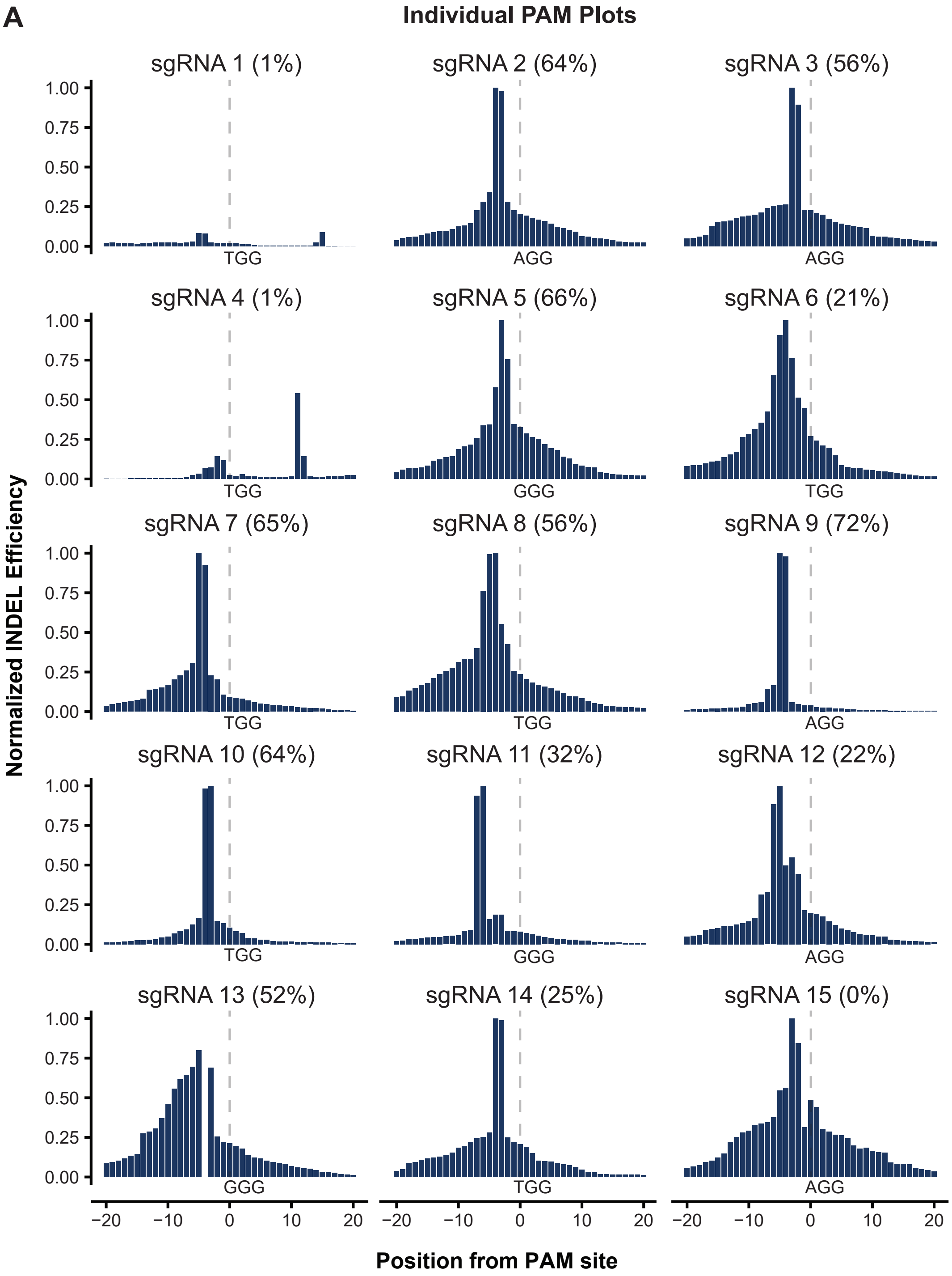


**Supplemental Figure 2. INDEL position relative to the OpenCRISPR-1 PAM sequences**

A) Normalized distribution of INDELs relative to OpenCRISPR-1 PAM sequence for each *Fut8* sgRNA screened.
