## Supplemental Table 1 for "An Efficient Workflow for CHO Cell Genome Engineering with OpenCRISPR-1"

| Target - Guide Name | Guide Sequence (5'-3') |
|---|---|
| Ces1f-1 | TCACCACTGGAGGTGAGGAT |
| Ces1f-2 | TGAACAGTGTTCACCACTGG |
| Ces1f-3 | CTGTTCATGGCAAAGTCCTG |
| Ces1f-4 | CTGTTCATGGCAAAGTCCTG |
| Ces1f-5 | CAAGAGGAGGCTTGGCAAAG |
| Ces1f-6 | AGCCTCCTCTTGGATCCCTG |
| Ces1f-7 | AACCTCAGGGATCCAAGAGG |
| Ces1f-8 | GCAAACCTCAGGGATCCAAG |
| Ces1f-9 | CTGTGGTGGAGCAAACCTCA |
| Ces1f-10 | GCATTCTTCACGAAGCTCCA |
| Cmah-1 | TTGGTTAATTGGCCCTGCTT |
| Cmah-2 | AAAGCAGGGCCAATTAACCA |
| Cmah-3 | GCATGTGACTGATGTAAATG |
| Cmah-4 | CAAGTTGGGAGACAAGCGGA |
| Cmah-5 | ACCTCAAGTTGGGAGACAAG |
| Cmah-6 | CTCTCCAGCCAATCAGATGG |
| Cmah-7 | ACCATCCTCGGGCAAAAGCA |
| Cmah-8 | CTGCTTTTGCCCGAGGATGG |
| Cmah-9 | GCATGGACCTCAAGTTGGGA |
| Cmah-10 | ATGTAGCAACCACCATCCTC |
| Fut8-1 | TAACAAAGAAGGGTCATCAG |
| Fut8-2 | TGACCCTTCTTTGTTAAAGG |
| Fut8-3 | CTTTGTCAGTGCGTCTGACA |
| Fut8-4 | GTGTACCATGTATTCCTCAA |
| Fut8-5 | CCATGTATTCCTCAATGGGA |
| Fut8-6 | CCATCCCATTGAGGAATACA |
| Fut8-7 | TTGCCTCCTTTAACAAAGAA |
| Fut8-8 | TTTGCCTCCTTTAACAAAGA |
| Fut8 Sun (2015)-1 | TGATGACCCTTCTTTGTTAA |
| Fut8 Sun (2015)-2 | AGCAGCCTTCCATCCCATTG |
| Fut8 Sun (2015)-3 | TGTACCATGTATTCCTCAAT |
| Fut8 Sun (2015)-4 | GTATTCCTCAATGGGATGGA |
| Fut8 Sun (2015)-5 | TCTCGAACGCAGAATGAAAG |
| Fut8 Wang (2018)-2 | GTCAGACGCACTGACAAAGT |
| Fut8 Wang (2018)-3 | GGATAAAAAAAGAGTGTATC |
| Ggta1-1 | TGACATAAAATATGACCCTG |
| Ggta1-2 | GTGTATAGAAGGCATTCTGG |
| Ggta1-3 | AACACTTGTAAGGAATGCAG |
| Ggta1-4 | GATGCGGATGAAGACCATCG |
| Ggta1-5 | CCTGTCCCACATACAGCACG |
| Ggta1-6 | GTGACAAGTTTACCTATGAG |
| Ggta1-7 | ACAAGTTTACCTATGAGAGG |
| Ggta1-8 | TGAGAGGCGGAAACTGTCGG |
| Ggta1-9 | TTCCTCCAAAAATGGCTGCG |
| Ggta1-10 | GCAATGACATCGAAGCAGAG |
| Lipa-1 | GATGCAGATCCTCGGCCTGG |
| Lipa-2 | AAAGACAGACCACCAGGCCG |
| Lipa-3 | TTTCGGTCCTGCTTTCTGGG |
| Lipa-4 | TTCGGTCCTGCTTTCTGGGA |
| Lipa-5 | TGCTTTCTGGGAGGCCCACG |
| Lipa-6 | CACGGGGTCCATTCCACATG |
| Lipa-7 | ACGGGGTCCATTCCACATGT |
| Lipa-8 | CGGGGTCCATTCCACATGTG |
| Lipa-9 | TTCCACATGTGGACCCCGAA |
| Lipa-10 | CATGTGGACCCCGAAGCAAA |
| LOC100768845-1 | TAAAACTCAAGTTTGAGCAA |
| LOC100768845-2 | CTCAAACTTGAGTTTTAGAG |
| LOC100768845-3 | AGGCTTGCCAAAGAGAAATG |
| LOC100768845-4 | AAAGAGAAATGTGGAAAACC |
| LOC100768845-5 | GAGAAATGTGGAAAACCAGG |
| LOC100768845-6 | TCAACATAATCACTAAGACT |
| LOC100768845-7 | TAATCACTAAGACTCGGCCA |
| LOC100768845-8 | GACTCGGCCAGGGACAGAGA |
| LOC100768845-9 | AGGGACAGAGAAGGCTAAGC |
| LOC100768845-10 | GGCTAAGCAGGAAATGAGTG |

| Target - Guide Name | Guide Sequence (5'-3') |
|---|---|
| LOC100769250-1 | CAGGAACACACACACAGGCC |
| LOC100769250-2 | CAACTGAGTGACACCAAAGT |
| LOC100769250-3 | GCTTGGCAAAGGGAATTCCC |
| LOC100769250-4 | CCCTTTGCCAAGCCACCTGT |
| LOC100769250-5 | AAGCGCAGTGGTCCGACAGG |
| LOC100769250-6 | TCAGGAGGTGCAAAGCGCAG |
| LOC100769250-7 | CTCCTGAAGCCCCTGAACCA |
| LOC100769250-8 | CATGGTTCAGGGGCTTCAGG |
| LOC100769250-9 | GAAGCCCCTGAACCATGGAG |
| LOC100769250-10 | CCATCTCTCACACCACTCCA |
| Lpl-1 | GAGCAAAGCCCTGCTCCTGG |
| Lpl-2 | TCCTGGTGGCTCTGGGAGTG |
| Lpl-3 | GGTGGCTCTGGGAGTGTGGC |
| Lpl-4 | CAGAGTTTGACCGCCTCCCA |
| Lpl-5 | AGTTTGACCGCCTCCCAAGG |
| Lpl-6 | GTTTGACCGCCTCCCAAGGA |
| Lpl-7 | TTTGACCGCCTCCCAAGGAG |
| Lpl-8 | GACCGCCTCCCAAGGAGGGG |
| Lpl-9 | GGCCACCCCTCCTTGGGAGG |
| Lpl-10 | TGCGGCCACCCCTCCTTGGG |
| Pla2g15-1 | TCACCTCACTTGTCGCGCGA |
| Pla2g15-2 | CACTTGTCGCGCGACCCAGC |
| Pla2g15-3 | CGCGCGACCCAGCTCCGGAG |
| Pla2g15-4 | CAAGAGGCCACTCCGGAGCT |
| Pla2g15-5 | CTTGGTCCCGCTGCTTCTGC |
| Pla2g15-6 | GCTGCTTCTGCTAATGATGC |
| Pla2g15-7 | CTGCTTCTGCTAATGATGCT |
| Pla2g15-8 | GCTTCTGCTAATGATGCTGG |
| Pla2g15-9 | CTCGGTCCAACGTCACCCCC |
| Pla2g15-10 | GGTCCAACGTCACCCCCCGG |
| Plbd2-1 | CAGCAACACCTCAGTCAGCG |
| Plbd2-2 | GGTGTTGCTGAATTGCCCGG |
| Plbd2-3 | CGGCGTCCAGCAGCACCGAG |
| Plbd2-4 | GGACGGCATCCATCCCTACG |
| Plbd2-5 | GGCGCTGGCCTCGCTGACTG |
| Plbd2-6 | GTCGGGTCAGCTGCGCCTGG |
| Plbd2-7 | CAGGCGCAGCTGACCCGACG |
| Plbd2-8 | CGGCATCCATCCCTACGCGG |
| Plbd2-9 | GCGCCAAAACCTCGACCCGC |
| Plbd2-10 | TGAGGTGTTGCTGAATTGCC |
| Plbd2 Zhang (2020) | GCCCCCATGGACCGGAGCCC |
| Ppt1-1 | GGTGGCTCCTAGCTGTGAGC |
| Ppt1-2 | AGGGCAGAAGGCTCACAGCT |
| Ppt1-3 | GACCAGGCGGCGCAGCACCA |
| Ppt1-4 | CAGATGTCCCAGCGACCAGG |
| Ppt1-5 | GTTCAGATGTCCCAGCGACC |
| Ppt1-6 | CCAGCGGTGTCAGCGAAGGC |
| Ppt1-7 | ACCAGCGGTGTCAGCGAAGG |
| Ppt1-8 | ATCACCAGCGGTGTCAGCGA |
| Ppt1-9 | CACCGCTGGTGATCTGGCAT |
| Ppt1-10 | ATCCCATGCCAGATCACCAG |
