## Supplemental Table 2 for "An Efficient Workflow for CHO Cell Genome Engineering with OpenCRISPR-1"

| Primer | Sequence (5'-3') |
|---|---|
| Ces1f | AGGGAAACCCACATGGAATC |
| Ces1f | TCCCACCTGCATCACTCTA |
| Cmah | TCGTCGGCAGCGTCAGATGTGTATAAGAGACAGGAGCCATCACTCCATTGTGGTTGAC |
| Cmah | GTCTCGTGGGCTCGGAGATGTGTATAAGAGACAGCACTGGTTATCTTAGGAGTTGGGCTTGG |
| Fut8 | ATCAGTGTGCTCTCCACTTC |
| Fut8 | TGCATCAGATACAGCTAGTAAGG |
| Fut8 Clones | TCGTCGGCAGCGTCAGATGTGTATAAGAGACAGGGGATACTAATTGAGTACCAGTACATTATCAGTGTG |
| Fut8 Clones | GTCTCGTGGGCTCGGAGATGTGTATAAGAGACAGCCCAGAATTAAGGATTTTCTTCAACTAAGAGATAATCCC |
| Ggta1-1 | GCATGTGTCCCCGTAAGGGTACTAG |
| Ggta1-1 | CAAACTATACTCTTTTGTCTGCCAGGC |
| Ggta1-1 Nested | TCGTCGGCAGCGTCAGATGTGTATAAGAGACAGGAGCATTATTTGGAAGATTTTCTGGAGTCTGC |
| Ggta1-1 Nested | GTCTCGTGGGCTCGGAGATGTGTATAAGAGACAGCCAGAGTCTCCACACCGAAATTGTC |
| Ggta1-2 | TCGTCGGCAGCGTCAGATGTGTATAAGAGACAGCCAGTTGGTAGCTCAGCTCCAGG |
| Ggta1-2 | GTCTCGTGGGCTCGGAGATGTGTATAAGAGACAGCCCAACAGTATTCTGGAGATAAGATTTTAGTGGG |
| Lipa | CTGTCTAAGTTCCTCTCCATGTC |
| Lipa | AGCACTTGTCATTGCTGTATTT |
| LOC100769250 | TCGTCGGCAGCGTCAGATGTGTATAAGAGACAGGGCATCTCTCCACACAGGTCAGG |
| LOC100769250 | GTCTCGTGGGCTCGGAGATGTGTATAAGAGACAGCTTCAGCATTGCTTAGAGCTCACAGC |
| Lpl | CCTTGCTACACGTATCTGCTC |
| Lpl | CTGAGTCTTTCCTTGCTCCTAC |
| Pla2g15 | TCGTCGGCAGCGTCAGATGTGTATAAGAGACAGCTAAAGCAGTCCATTGGACCTGCTG |
| Pla2g15 | GTCTCGTGGGCTCGGAGATGTGTATAAGAGACAGGACCCGACCCAGGTTAAGAGACC |
| Plbd2 | TCGTCGGCAGCGTCAGATGTGTATAAGAGACAGGCACGCGATGGCGGC |
| Plbd2 | GTCTCGTGGGCTCGGAGATGTGTATAAGAGACAGCGGCCAACCCTTACCCGG |
| Ppt1 | TCGTCGGCAGCGTCAGATGTGTATAAGAGACAGGATGGCGTCGCCCGGC |
| Ppt1 | GTCTCGTGGGCTCGGAGATGTGTATAAGAGACAGGTGCACAGCTCTCCCCGAGAG |
